# Respiratory gas levels interact to control ventilatory motor patterns in isolated locust ganglia

**DOI:** 10.1101/543793

**Authors:** Stav Talal, Amir Ayali, Eran Gefen

**Author notes:** Corresponding author **Corresponding author’s**.

## Abstract

Large insects actively ventilate their tracheal system even at rest, using abdominal pumping movements, which are controlled by a central pattern generator (CPG) in the thoracic ganglia. We studied the effects of respiratory gases on the ventilatory rhythm by isolating the thoracic ganglia and perfusing its main tracheae with various respiratory gas mixtures. Fictive ventilation activity was recorded from motor nerves controlling spiracular and abdominal ventilatory muscles. Both hypoxia and hypercapnia increased the ventilation rate, with the latter being much more potent. Sub-threshold hypoxic and hypercapnic levels were still able to modulate the rhythm as a result of interactions between the effects of the two respiratory gases. Additionally, changing the oxygen levels in the bathing saline affected ventilation rate, suggesting a modulatory role for haemolymph oxygen. Central sensing of both respiratory gases as well as interactions of their effects on the motor output of the ventilatory CPG reported here indicate convergent evolution of respiratory control among terrestrial animals of distant taxa.

**Summary statement:** Tight control over respiratory gas supply to the isolated locust CNS reveals interactions of oxygen and carbon dioxide effects on central ventilatory output.

## Introduction

Insects use a system of gas-filled trachea to exchange respiratory gases with their external environment. Large insects actively ventilate their tracheal system in order to satisfy their respiratory demands even at rest (Harrison, 1997). The best studied models for respiration and ventilation regulation are locusts and cockroaches (e.g. Burrows, 1996; Harrison, 1997; Harrison et al., 2013; Matthews, 2017; Miller, 1966). Under increased metabolic demands these insects increase their respiratory gas delivery by increasing their tidal volume by way of recruiting auxiliary respiration muscles (Miller, 1960; Weis-Fogh, 1967) and a higher ventilation rate (reviewed in Harrison, 1997; Matthews, 2017; Miller, 1966). Numerous studies have shown that resting insects increase ventilation rate in response to an exposure to experimental hypercapnic/hypoxic conditions (e.g. Gulinson and Harrison, 1996; Krolikowski and Harrison, 1996; Matthews and White, 2011). Studies of the chemosensory mechanisms by which ventilation is controlled, have been mainly conducted on intact insects, using manipulation of external gas levels or, alternatively, manipulation of endo-tracheal respiratory gases by means of tracheal system perfusion (e.g. Arieli and Lehrer, 1988; Groenewald et al., 2014; Gulinson and Harrison, 1996; Huang et al., 2014; Hustert, 1975; Matthews and White, 2011; Talal et al., 2018). Hence, the physiological factors that have been suggested to regulate the ventilation rate include the endo-tracheal and haemolymph respiratory gas levels (Gulinson and Harrison, 1996; Miller, 1966, also reviewed in Matthews, 2017; Matthews and Terblanche, 2015), and the haemolymph pH (Farley and Case, 1968; Snyder et al., 1980).

The neural control of the ventilation rhythm in insects has been extensively studied (reviewed by Burrows, 1996; Miller, 1966; Miller, 1981). In locusts and cockroaches, the source of the ventilation pattern was reported to be an endogenous pacemaker or a central pattern generator (CPG) circuit located in the metathoracic ganglion complex (comprising in locusts the metathoracic ganglion fused with the first three abdominal ganglia; reviewed in Burrows, 1996). In locusts, direct CO_2_ perfusion of head and thoracic ganglia in an exposed neural system preparation resulted in increased ventilation rate (Miller, 1960). Moreover, several members of the ventilatory CPG, i.e. interneurons whose activity could change the rhythmic pattern, have been identified in the metathoracic ganglion (Pearson, 1980; Ramirez and Pearson, 1989). Bustami and colleagues (Bustami and Hustert, 2000; Bustami et al., 2002) were the first to demonstrate changes in the rate of fictive (*in-vitro*) ventilation using an isolated CNS preparation comprised of the thoracic ganglia and the first unfused abdominal ganglion. These studies demonstrated a change in the output of the CPG in response to manipulation of the respiratory gas levels in the gas phase above the saline in which the preparation was bathed. The authors suggested the existence of oxygen and CO_2_/pH sensors that modulate the rate of the ventilatory CPG (Bustami et al., 2002). However, despite their important pioneering work, some of the findings were at odds with observations from intact insects. For example, the effect of hypercapnia on the fictive ventilation rate was recorded at 20% CO_2_ (Bustami et al., 2002), a considerably higher level than those eliciting hyperventilation in experiments using intact insects (2-3%; Gulinson and Harrison, 1996; Harrison et al., 1995; Hustert, 1975; Matthews and White, 2011). Moreover, the PCO_2_ level measured in metathoracic trachea was only ~2% in postexercise grasshoppers which exhibited a 6-8 fold increase in ventilation rate (Table 1 and 2 in Krolikowski and Harrison, 1996).

Modulation of ventilatory motor patterns is well exemplified in insect discontinuous gas exchange (DGE). This most studied insect gas exchange pattern is exhibited during periods of quiescence or at low metabolic rates (reviewed in Chown, 2011; Chown et al., 2006; Contreras and Bradley, 2009; Lighton, 1996; Matthews, 2017; Terblanche and Woods, 2018). It comprises three phases, defined by the spiracle state as reflected by CO_2_ emission rate: close, flutter and open. The transition between the DGE phases is triggered by chemoreceptors that respond to changes in internal respiratory gas levels. It is well established that when insects employ DGE, a build-up in internal CO_2_ levels triggers spiracle opening and a bout of rapid gas exchange (Förster and Hetz, 2010; Levy and Schneiderman, 1966a). A decrease in tissue oxygenation level was also shown to trigger ventilatory movements in order to mix tracheal gaseous contents when spiracles are closed (Huang et al., 2014). Several recent studies focused on the DGE pattern in actively ventilating insects such as locusts and cockroaches. Ventilation was shown to be tightly coupled to the spiracle activity, creating a unidirectional air flow through the body (Heinrich et al., 2013; Matthews and White, 2011; Talal et al., 2018). It was further demonstrated that the rate of ventilation changes continuously during the open phase (during the active ventilation of the tracheal system in order to exchange respiratory gases), probably in response to changes in the internal gas levels (Matthews and White, 2011; Talal et al., 2018). A number of studies have reported that the duration of the closed phase is extended in response to hyperoxic conditions, even when metabolic rates remained unchanged, thus suggesting interactions between the effects of the different respiratory gases in the regulation of DGE (Levy and Schneiderman, 1966b; Matthews and White, 2011; Talal et al., 2018). Periodic fictive ventilation events, reminiscent of DGE were demonstrated in the isolated locust thoracic ganglia preparation mentioned above (Bustami and Hustert, 2000).

The current study was therefore aimed at characterizing tentative interactions between the modulatory effects of the respiratory gases and their potential role in the control of ventilation and gas exchange motor patterns in insects. We hypothesized that ventilatory regulation is sensitive to buildup of CO_2_ in the tracheal system, as it is to interacting effects of internal O_2_ and CO_2_ levels as observed in intact insects (Levy and Schneiderman, 1966b; Matthews and White, 2011; Talal et al., 2018), and indeed in terrestrial animals of various taxa, including mammals (reviewed in Teppema and Dahan, 2010). In order to achieve full control over the experimental conditions, and to validate the physiological relevance of previous reports, we utilized an isolated *in vitro* preparation of the locust thoracic ganglia, together with their associated ventral longitudinal tracheal trunks. This allowed delivery of tightly controlled respiratory gas mixtures of various compositions to the isolated ganglia, while simultaneously controlling saline pH and gas levels. Ventilatory motor patterns were continuously monitored by recording efferent discharges under the different experimental conditions.

## Materials and Methods

### Experimental insects

In this study we used desert locusts *(Schistocerca gregaria)* from our stock population at the Tel Aviv University. These were kept at 33.0±3.0°C, under a 14L:10D photoperiod (supplementary radiant heat was supplied during daytime by incandescent 25W electric bulbs). Locusts were fed daily with wheat shoots and dry oats *ad libitum.* All experiments were carried out on males only, 1-3 weeks after adult eclosion.

### In-vitro preparation

The locusts were anesthetized on ice prior to dissection. Following the removal of legs, wings, the pronotum and the abdomen posteriorly to the fourth abdominal segment, a longitudinal cut was performed in the cuticle along the dorsal midline of the thorax. The preparation was attached to a Sylgard dish (Sylgard 182 silicon Elastomer, Dow Corning Corp., Midland, MI, USA), the cut was widened and the gut removed. The body cavity was perfused with locust saline (Knebel et al., 2019). Air sacs and fatty tissue covering the ventral nerve cord were removed. The thoracic ganglia chain, comprising the three thoracic ganglia (pro-, meso- and meta-thoracic ganglion) and the first unfused abdominal ganglion with the surrounding tracheal supply were dissected out, pinned in a clean Sylgard dish, dorsal side up, and bathed in locust saline. The two main tracheae were opened from both sides and floated on the saline surface (Fig. 1).

**Figure 1.**
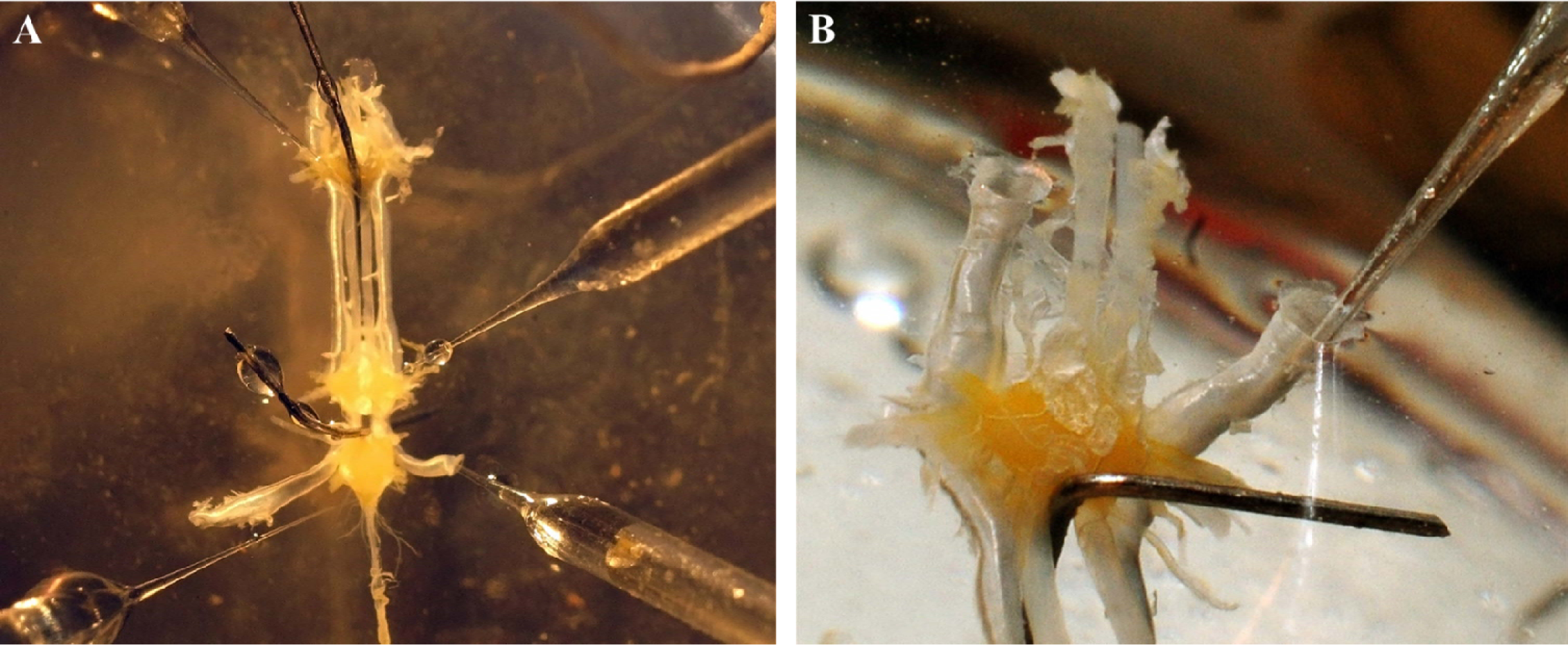
The *in-vitro* preparation. **A**. The three (pro-, meso- and meta-) thoracic ganglia bathed in saline with the pair of ventral longitudinal tracheae. Custom-made suction electrodes were used to record fictive ventilation discharge from the nerve stumps. An additional glass capillary (lower right) was inserted into one end of the ventral longitudinal trachea in order to manipulate the endo-tracheal respiratory gas content. **B**. An enlarged view of the pro-thoracic ganglion (different preparation) with the two ventral longitudinal tracheae and the gas-manipulating glass capillary prior to saline submergence.

### Tracheal and saline perfusion, and respiratory gases manipulation

Unless stated otherwise, saline was bubbled with pure nitrogen at a flow rate of 25 ml·min^-1^ using a mass-flow controller (FMA-2617A; Omega Engineering, Inc., Stamford, CT, USA). This created an anoxic saline (verified using an IMP-PSt1 PO_2_ optode and a Microx TX3, micro fiber oxygen transmitter, PreSens, Regensburg, Germany) in order to prevent oxygen diffusion to the nervous system, and to ensure that tissue oxygenation was limited to tracheal supply only.

We manipulated CO_2_ and O_2_ levels in the main longitudinal tracheae, which supply the thoracic ganglia with respiratory gases (Burrows, 1980; Fig. 1B). Respiratory gas mixtures were perfused through a glass capillary directly into the tracheae at a flow rate of 5 ml·min^-1^. We used certified gas mixtures (79% N_2_/ 21% O_2_; 87% N_2_/ 7% CO_2_/ 6% O_2_; 87.5% N_2_/ 3.5% CO_2_/ 9% O_2_; 94% N_2_/ 6% O_2_; pure N_2_) and two mass-flow controllers (FMA-2616A; Omega Engineering Inc., Stamford, CT, USA) to achieve the desired experimental gas compositions. At an accuracy/repeatability of 0.2% of full scale (20 ml·min^-1^ for the FMA-2616A) the error was <0.2% and 0.1% for O_2_ and CO_2_ concentrations, respectively. Following the glass capillary insertion (usually at the anterior end near the prothoracic ganglion) (Fig. 1B), the tracheal opening was pushed under the saline resulting in collapse of the tracheal walls against the capillary tube and its sealing. In order to ensure perfusion of the gas mixtures through both ventral tracheae (which are connected through air sacs under each thoracic ganglion) and to make sure that the perfused gases leave the tracheal system only from the far end, the tracheal opening contralateral to the capillary-connected one was also pushed under the saline surface and collapsed shut. The preparation temperature was monitored and maintained at 24±1°C.

### Extracellular recording from nerve stumps

We used custom-made suction electrodes to record extracellularly the ventilation motor pattern activity from the median nerves of the prothoracic and metathoracic ganglia, which innervate the first thoracic spiracle closer muscle and the ventilatory and spiracle closer muscles of the 4^th^ abdominal segment, respectively (Fig. 1A). Data were acquired and stored on a computer for offline analysis using a four-channel differential amplifier (Model 1700, A-M Systems, Sequim, WA, USA) and Axon Digidata 1440A A-D board with Axo-Scope software (Molecular Devices, Sunnyvale, CA, USA).

### Experiments

Each experiment started 30 minutes following the insertion of the electrodes and the gas-perfusion capillary. Each preparation was exposed to “control” gaseous conditions (see below) for 30 minutes before tracheal gas composition was changed for 30 minutes, followed by an additional 30 minutes exposure to the initial (control) gas composition. Endo-tracheal oxygen or carbon dioxide levels were first manipulated separately while the other gas was kept constant. Next, several additional gas level combinations were tested to elucidate the interaction between tracheal levels of the two respiratory gases. The initial and final exposures to control conditions served to verify the tissue viability and rule out any potential effects of the long exposure to dry tracheal contents.

In order to rule out the possibility that our findings reflected tissue response to anoxic saline conditions, we compared fictive ventilation rates in anoxic saline to those recorded from preparations that were bathed in hypoxic (6% O_2_ in N_2_) and normoxic (21% O_2_ in N_2_) saline (achieved by bubbling the respective gas mixes into the saline). These experiments were carried out while perfusing the trachea with 6% O_2_ and 3.5% CO_2_ (in N_2_).

### Data analysis and statistics

Extracellular recordings were analysed using DataView 10.6 (W.J. Heitler, University of St. Andrews, Scotland). Bursts of action potentials were identified and their rate (the fictive ventilation rhythm frequency) was determined from the last 10 min of each 30 min data recording session (no unit identification or sorting was performed). Following normality tests (Kolmogorov-Smirnov and Shapiro-Wilk), we used paired t-test to compare between the fictive ventilation rates at the manipulated and control respiratory gas levels. Log_2_ transformation was applied to the one data set that failed the normality tests, allowing the use of the paired t-test as above for this case also. We compared the effect of different saline oxygen levels on the ventilation rate using one-way ANOVA, followed by Bonferroni post-hoc test. Statistical analyses were performed using SPSS 20.0 (IBM).

## Results

Simultaneous extra-cellular recordings of the median nerves of the prothoracic ganglion and the third fused abdominal ganglion in our *in-vitro* preparation revealed robust fictive rhythmic ventilation-related motor patterns. The recorded pattern was characterized by perfect burst alternation (Fig. 2) that in the intact insect would result in alternations between activity of the anterior and posterior spiracles’ muscles and the ventilatory muscles. The *in-vitro* frequency recorded in our control conditions was ~5 bursts·min^-1^ at 24±1 °C.

**Figure 2.**
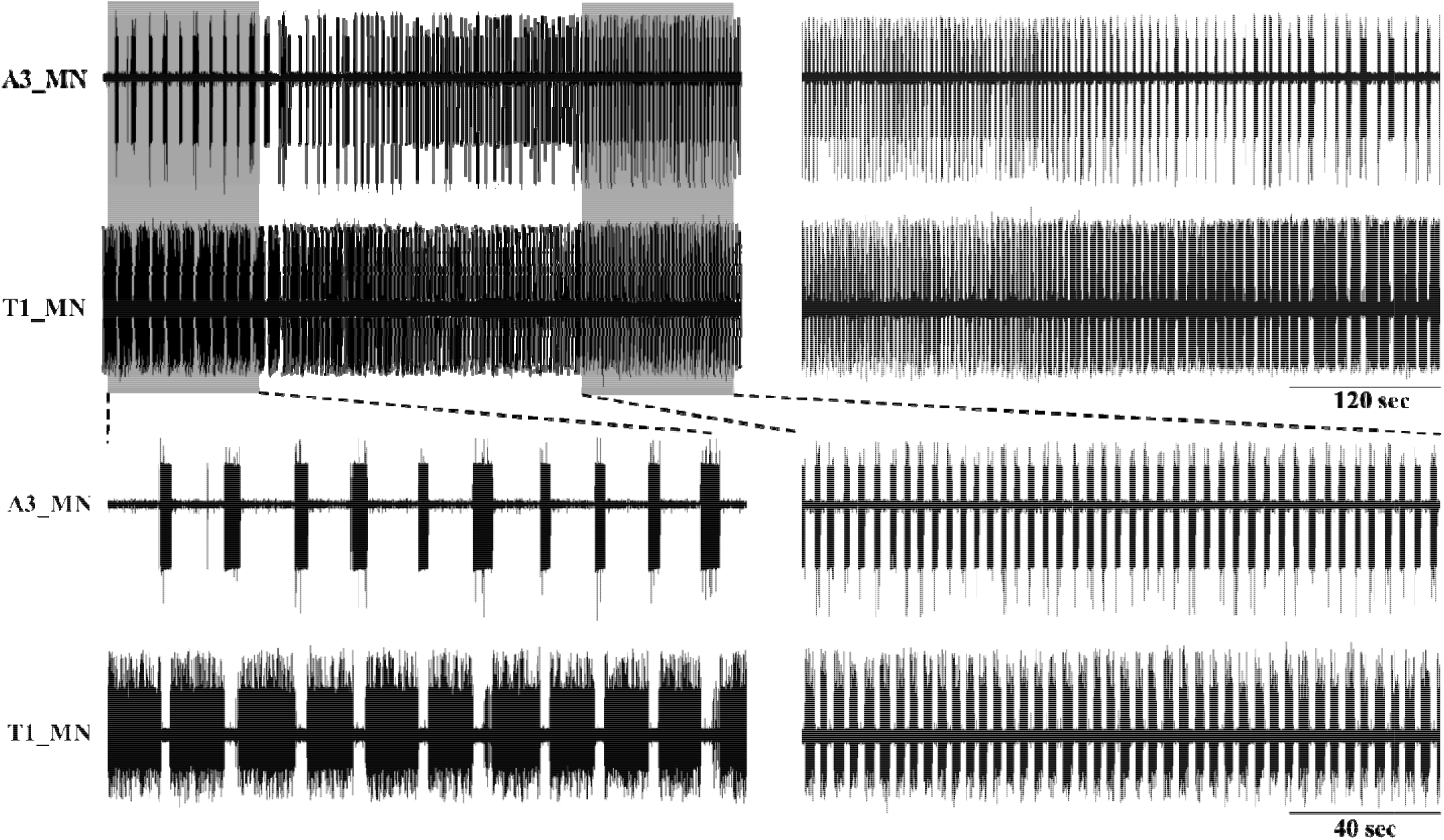
An example of the time course of the change in the fictive ventilation rate in response to elevated endo-tracheal CO_2_ level (5%) (upper left) and return to the initial 0% CO_2_ (upper right), with endo-tracheal oxygen level remaining constant at 6%. The shaded areas are enlarged in the lower panel. A3_MN and T1_MN denote the median nerves of the third abdominal ganglion (which is fused with the meta-thoracic ganglion) and the pro-thoracic ganglion, respectively.

Although we used only the last 10 minutes of each 30 minute exposure to the experimental conditions for our data analysis, changes in the efferent discharge in response to changing tracheal gas levels were typically evident within 2-5 minutes (Fig. 2). Decreasing tracheal oxygen level to 2% resulted in a small but significant increase in the fictive ventilation rate (Fig. 3A; t_6_=5.702, p=0.001; CO_2_ kept at 0%). No response was recorded however at 10% or 4% O_2_ (t6=0.672, p=0.527 and t5=0.882, p=0.418, respectively).

**Figure 3.**
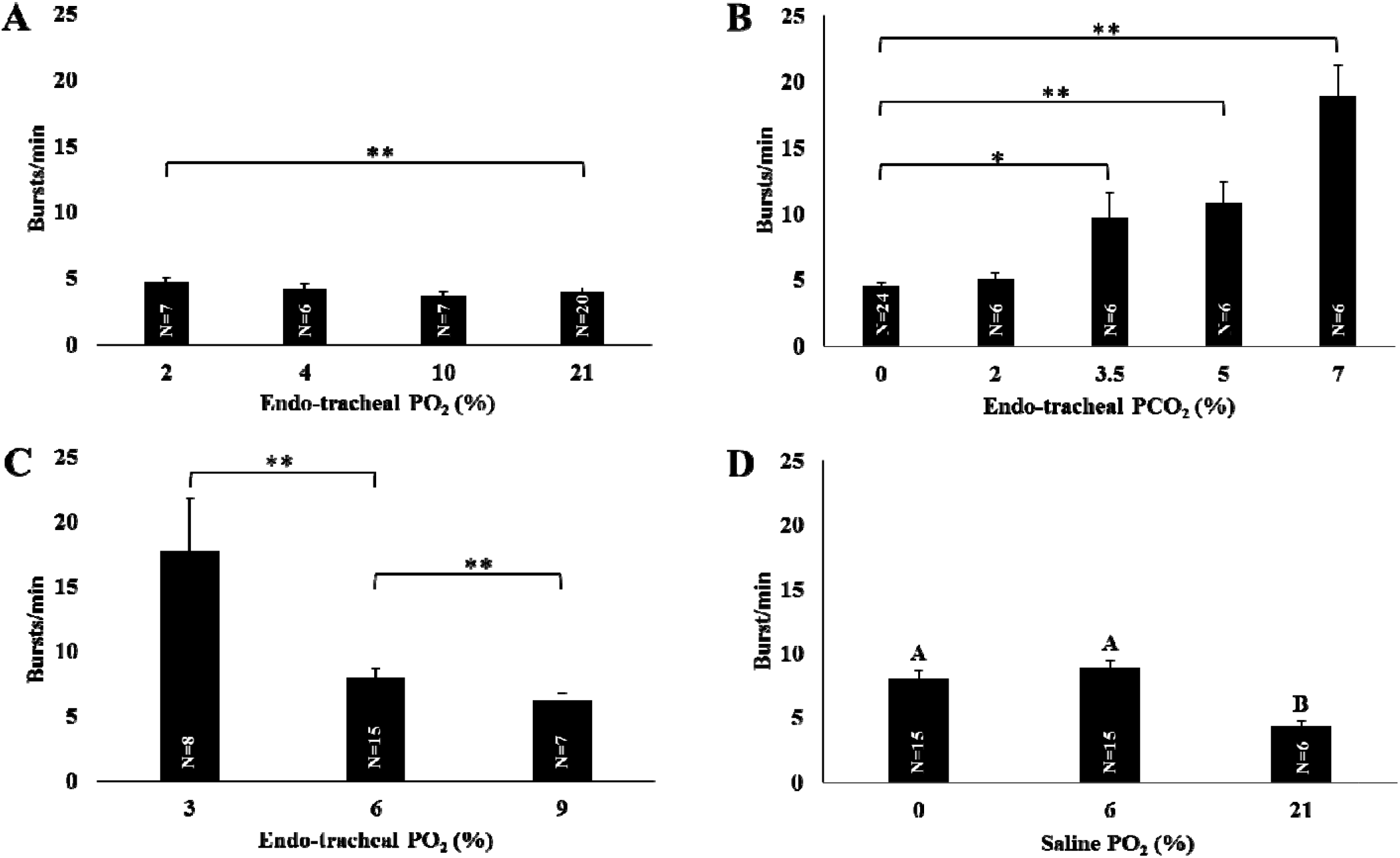
Changes in the fictive ventilation rate in response to different endo-tracheal/saline respiratory gas levels: **A**. Effect of endo-tracheal O_2_ level (%, at 0% CO_2_). **B**. Effect of endotracheal CO_2_ level (%, at 6% O_2_). **C**. Effect of endo-tracheal O_2_ level (%, at 3.5% CO_2_). **D**. Effect of saline oxygenation level (%, when endo-tracheal gas concentrations were kept constant at 6% O_2_ and 3.5% CO_2_). Asterisks indicate significant differences: *p<0.05; **p<0.01. Values are means ± s.e.m.

Preliminary experiments in which the tracheal perfusion comprised 21% O_2_ did not result in a response to elevated (up to 7%) CO_2_ levels (data not shown). Consequently, we studied the effect of tracheal CO_2_ on the ventilatory pattern while maintaining the tracheal O_2_ level at 6% (which was above the recorded oxygen threshold for inducing increased ventilation rate; Fig. 3A), similar to haemolymph values measured in a continuously ventilating insect (P.G.D. Matthews, pers. comm). Overall, the effect of changing the CO_2_ levels was much more pronounced compared to the relatively limited O_2_ effect (Fig. 3B). The threshold for the CO_2_ effect was found to be between 2% and 3.5%. A two-fold increase in the fictive ventilation rate was recorded at 3.5% and at 5% CO_2_ (t_5_=2.582, p=0.049; t_5_=4.614, p=0.006, respectively), while 7% CO_2_ resulted in a four-fold increase (t_5_=6.558, p=0.001). As noted, there was no increase in the ventilation rate when the endo-tracheal CO_2_ level was 2% (t_5_=0.999, p=0.364; Fig. 3B). Tissue viability during the duration of the experiment was confirmed by the almost identical burst frequencies at the same gaseous exposure at the beginning and end of each experiment (t_17_=0.002, p=0.99 and t_18_=0.65, p=0.53 for the O_2_ and the CO_2_ manipulation experiments, respectively).

While neither 2% CO_2_ nor O_2_ ≥4% elicited a change in the fictive ventilation rate when each gas was tested separately (Figs. 3A, 3B), an interaction between the effects of the two gases was evident when applying different gas level combinations. In a separate set of measurements (N=7), changing tracheal gas levels from 3.5% CO_2_ / 6% O_2_, to 2% CO_2_ / 2% O_2_, resulted in an increase in the ventilation rate from 6.4±0.8 to 15.4±2.3 bursts per minute (t_6_=3.385; p=0.015). Similarly, when tracheal CO_2_ level was maintained at 3.5% (above our recorded threshold for hyperventilation; Fig. 3B), changing the O_2_ level from 6% to 3% resulted in a two-fold increase in fictive ventilation rate (t_7_=3.656; p=0.008; following log_2_ transformation to reach normality; Fig. 3C), whereas a significant decrease was recorded at 9% O_2_ (t_6_=4.245; p=0.005; paired t-test; Fig. 3C).

Saline perfusion with nitrogen resulted in anoxic saline and prevented oxygen diffusion from the saline surface to the submerged ganglia. In order to rule out a possible artifact resulting from the anoxic saline in which the preparation was bathed, we carried out a complementary set of experiments in which we changed the saline oxygen level by means of perfusion of different gas mixtures (pure N_2_, 6% O_2_ in N_2_ and 21% oxygen in N_2_), while the main trachea was constantly perfused with 6% O_2_ and 3.5% CO_2_. We found that the saline oxygen level indeed affected the ventilation rate (Fig. 3D). Under normoxic saline the ventilation rate was significantly lower compared to hypoxic and anoxic saline (F_2,33_=7.57; p=0.002 one-way ANOVA; Fig. 3D). Furthermore, 3.5% CO_2_ failed to elicit hyperventilation (rates were no different from those for 0% CO_2_; Fig. 3B).

## Discussion

The existence and even the location of the ventilatory CPG in insects are known for more than half a century (reviewed in Ayali and Lange, 2010; Burrows, 1996; Miller, 1966). However, our knowledge of the mechanisms related to gas sensing is still very limited. The findings by Bustami et al. (2002) provided the first direct evidence of the existence of oxygen chemoreceptors within the locust CNS. However, limited control over the precise gas levels close to the site of the respiratory CPG in their experiments obscured a significant response to certain physiologically-relevant gas conditions. We developed a novel *in-vitro* preparation that allowed us to examine the mechanisms of chemosensory modulation of the endogenic ventilatory CPG(s), while controlling the immediate respiratory gas environment. We achieved this by using direct tracheal perfusion of respiratory gas mixtures varying in composition, while controlling saline oxygenation level.

The robust and consistent alternation between the bursts of action potentials recorded from the median nerves of different ganglia throughout our experiments supports the notion that the generation and control of the rhythmic ventilation-related motor patterns are hard-wired in the insect central nervous system (i.e. a system of CPGs), and result in a functional unidirectional air flow in the intact insect. Furthermore, our findings clearly demonstrate the existence of CO_2_ and oxygen sensing mechanisms in the thoracic ganglia, and their ability to manipulate the fictive ventilation-related output of the CPG. Throughout our experiments there was a clear time-lag between a change in the perfused gas mixture and the observed change in ventilation rate (see Fig. 2). This lag was probably only a result of a combination of the flow rate and the length of the tubing used for supplying the respiratory gas mixtures.

The most interesting result in the current study is that of the interaction between the respiratory gases in their effect on the fictive ventilation rate. We have shown that the threshold level (or the sensitivity) for the modulatory effect of each gas depends on the level of the other. For example, the fictive ventilation rate doubled when tracheal oxygen level was decreased from 6% to 3% at 3.5% CO_2_, whereas at 0% CO_2_ a significant (and milder; by ~20%) response was recorded only when tracheal oxygen level was lowered to 2%. Moreover, ventilation rate at 2% O_2_ / 2% CO_2_ (15.4±2.3 bursts per minute) was three-fold higher than the rate when each gas was kept at 2% separately, while the other was at sub-(for CO_2_; Fig. 3A) or above (for O_2_; Fig. 3B) threshold values. Notwithstanding the low oxygen threshold level reported here (Fig. 3A), intact locusts hyperventilate when the ambient oxygen level is reduced to ~10% (Bustami et al., 2002; Greenlee and Harrison, 1998). This apparent difference between the *in-vitro* preparation and intact insects could be explained by the aforementioned interaction between modulatory effects of the two respiratory gases. A 2% oxygen threshold was recorded at 0% CO_2_ (Fig. 3A) whereas the endo-tracheal CO_2_ level of the intact insects is 2-5% (Gulinson and Harrison, 1996; Harrison et al., 1995; Levy and Schneiderman, 1966a). We show that at 3.5% CO_2_ a significant increase in fictive ventilation was recorded when tracheal oxygen was lowered from 9 to 6 % (Fig. 3C) to a rate two-fold higher than normoxic values (Fig. 3A). The combined effects of the respiratory gases have also been found in intact cockroaches, in which the ventilation rate increased significantly more when exposed simultaneously to hypoxia and hypercapnia compared to the exposure to the same O_2_/CO_2_ condition separately (Matthews and White, 2011).

The interacting effects of respiratory gases are also apparent when insects perform DGE. The CO_2_ threshold level for triggering the open phase is elevated from 5% to 8% in lepidopteran pupae exposed to hyperoxia (Levy and Schneiderman, 1966b). In addition, lower haemolymph pH levels (a proxy to higher haemolymph, and tracheal PCO_2_) were measured in cockroaches under hyperoxic conditions just prior to the initiation of the open phase (Matthews et al., 2011). Likewise, lower endo-tracheal oxygen levels triggered the flutter phase in locusts under various hypoxic levels (Matthews et al., 2012), possibly as a result of hypoxia-induced hyperventilation and hemolymph CO_2_ washout (Matthews and White, 2011). We also showed that locusts performing DGE exhibit longer interburst phase under hyperoxic conditions compared with controls, despite their comparable CO_2_ accumulation rates during spiracle closure, suggesting a higher CO_2_ threshold for the open phase triggering in hyperoxia (Talal et al., 2018). This synergistic interaction could hint at a potential cellular oxygen sensing mechanism which is mediated by CO_2_ and/or pH level.

Respiratory regulation by a cellular O_2_/CO_2_ sensing mechanism has never been studied in insects. In contrast, research progress has been made in other terrestrial animals, and mammals in particular. Findings in this study indicate a similar response in actively ventilating insects to that found in mammals (and other vertebrates), which increase their ventilation rate and/or tidal volume in response to local hypoxia or hypercapnia (Teppema and Dahan, 2010). In contrast to insects, tissue gas exchange in vertebrates is mediated by the circulatory system. Arterial blood oxygen, CO_2_ and pH levels are monitored by the glomus cells, located in the aortic and the carotid bodies, and whose output modulates the ventilation rate (reviewed in Costa et al., 2014; Kumar and Prabhakar, 2012). There is evidence for interactions between the effects of oxygen and CO_2_ on peripheral chemoreceptors in mammals (reviewed in Teppema and Dahan, 2010). Interestingly, the synergistic effect of tracheal hypoxia and hypercapnia on respiratory CPG in locusts reported here is of similar shape to the previously reported response of the cat carotid body to changing gas partial pressures in the arterial blood (Hornbein et al., 1961) (Fig. 4). Additionally, hyperoxic and hypocapnic perfusions of the carotid sinus in dogs inhibit carotid body activity (Blain et al., 2009). Reduced sensitivity of the locust CPG was also observed in hypocapnia (Fig. 3A) or hyperoxia (Fig. 3D; Bustami et al., 2002). Hence, it is likely that the basic mechanism underlying respiratory gas chemosensing in mammalian carotid and aortic bodies and in the insect ventilatory CPG are similar, despite their independent evolution.

**Figure 4.**
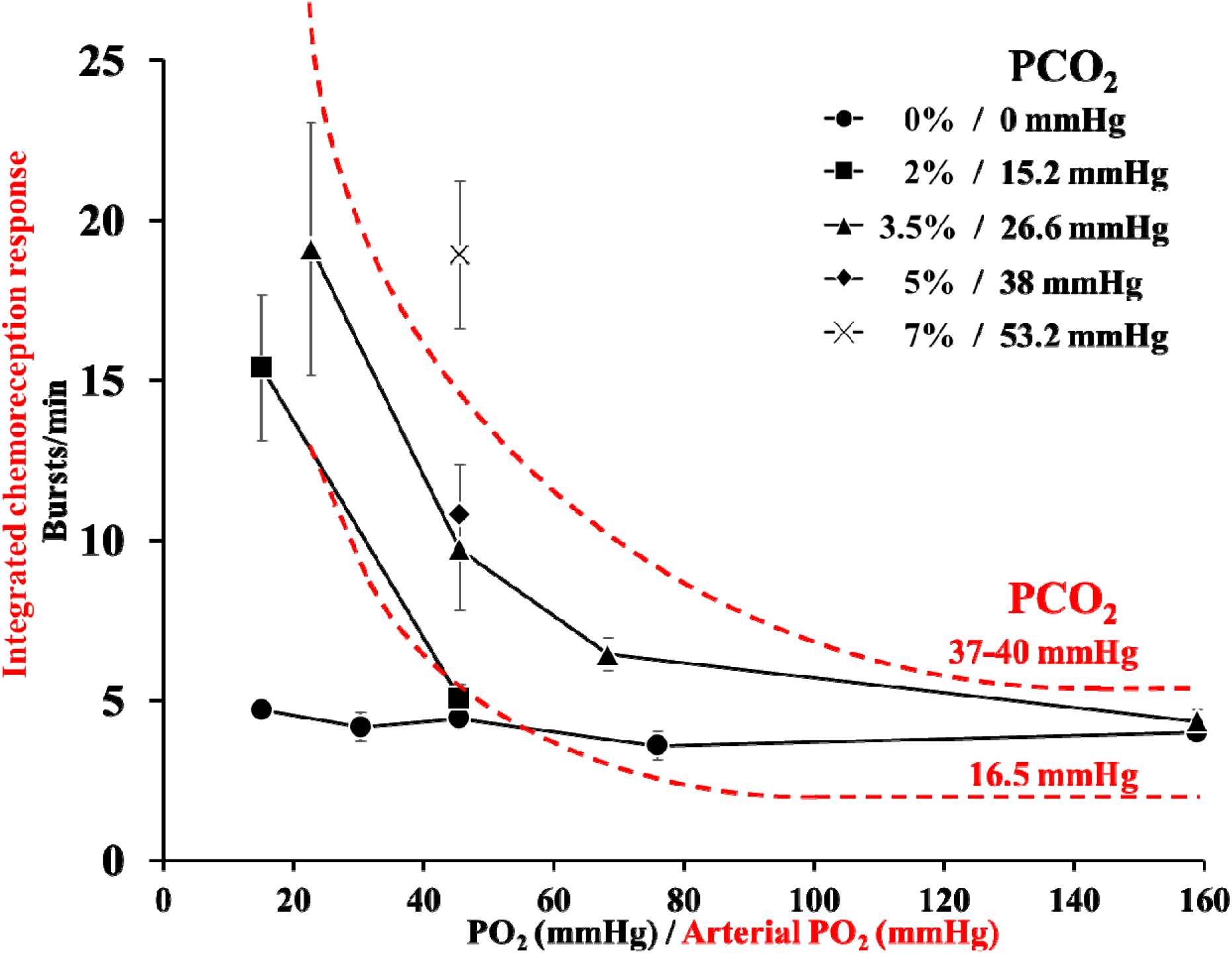
Chemosensory response to changing PO_2_ at different PCO_2_ levels in the cat carotid body (dashed red lines; modified from Hornbein et al., 1961; Fig. 4 therein) and the locust ventilatory CPG (Values are means ± s.e.m.).

Another novel insight offered by the current study is a role for hemolymph chemosensing in ventilatory CPG modulation. While the initial motivation for manipulating saline oxygenation level was to account for possible effects of anoxic saline, we were able to show that ventilatory response to tracheal level of 3.5% CO_2_ is maintained when saline O_2_ is increased from 0 to 6%, but abolished altogether in air-saturated saline (Fig. 3D). These manipulated O_2_ levels are within the PO_2_ range reported for the intact insect (4-10kPa in a flight muscle of resting hawkmoth, Komai, 1998; ~6.3±2.3kPa in the haemolymph of a lepidopteran pupa during continuous gas exchange, P.G.D. Matthews, pers. comm). This result could explain the previously reported high CO_2_ threshold (20%) in a study which used a thoracic ganglia preparation bathed in hyperoxic saline (50% O_2_; Bustami et al., 2002).

In conclusion, the current study provides important novel insights into insect central gas chemosensing and its control of ventilation. It also indicates an intriguing convergence with regulatory mechanisms in phylogenetically-distinct terrestrial vertebrates. However, the location, cellular mechanism and number of sensing sites are still unknown. The relatively tractable insect preparation, together with rapidly-developing genetic tools, could help in revealing the underpinning of respiratory gas sensing mechanism in insects and thus shed light on the vertebrate and even human systems, for which some controversies and gaps still exist.

## Authors’ contributions

E.G. and A.A. acquired the funding. All authors conceived and designed the experiments. S.T. performed the experiments and analyzed the data. All authors contributed to the manuscript preparation. All authors agree to be held accountable for the content therein and approve the final version of the manuscript.

## Acknowledgements

We thank Dr. Daniel Knebel and Izhak David for helping us with locust surgical procedures and with the electrophysiological setup. We also thank two anonymous reviewers for their constructive comments which helped improve a previous version of the manuscript.

